# Regulation of shelterin proteins TERF2IP and TRF2 by H3K4me3-p65 axis drives hyperglycemia dependent endothelial senescence

**DOI:** 10.1101/2023.05.12.540614

**Authors:** Sumukh Thakar, Ritobrata Bhattacharya, Yash T Katakia, Srinjoy Chakraborty, S Ramakrishnan, Niyati Pandya Thakkar, Smita Dey, Shibasish Chowdhury, Syamantak Majumder

## Abstract

**Background:** Endothelial senescence has been linked to several cardiovascular diseases. Dysregulation of proteins of the shelterin complex including TRF2 and TERF2IP causes senescence as it hampers DNA repair and cell proliferation. However, whether exposure to hyperglycemia interplays with proteins of the shelterin complex thus further dictates the senescent phenotype of endothelial cells (EC) remain to be explored.

**Approach and Results:** In this study, we observed elevated levels of p21 and p53 in endothelial cells upon exposure to intermittent hyperglycemia. We also noted hyperglycemia exposure increased the levels of TERF2IP and TRF2, part of the shelterin complex. No change in the level of TRF1 and TPP1 were detected. Furthermore, a robust induction was detected in p65 level upon intermittent hyperglycemia challenge. ChIP-qPCR analysis revealed enhanced H3K4me3 enrichment in the promoter regions of p65, TERF2IP and TRF2. Inhibition of catalysis of H3K4me3 either by pharmacological inhibitor or siRNA-mediated knockdown of MLL2 attenuated increase in p65, TERF2IP and TRF2 levels including reversal of senescence markers p53 and p21. Interestingly, pharmacological inhibition of NF-κB signaling also diminished abrupt increase in TERF2IP and TRF2 levels thus further reversed intermittent hyperglycemia-induced p53 and p21 levels. More importantly, co-immunoprecipitation and co-localization analysis revealed an interaction between nuclear p65 and MLL2 in EC stimulated with hyperglycemia. Further knockdown of either TERF2IP or TRF2 impaired intermittent hyperglycemia-induced p53 and p21 expression and associated endothelial senescence.

**Conclusion:** Overall, the present study describes an interplay of epigenetics in defining NF-κB signaling and shelterin proteins’ expression which further govern the biochemical and functional state of endothelial senescence in hyperglycemic milieu.

## Introduction

Diabetes-associated complications are one of the most common causes of cardiovascular and kidney diseases worldwide. Diabetes-associated complications are mainly manifested by both microvascular and macrovascular anomalies. Endothelial cells (EC) being the inner lining of the vasculature play a major role in maintaining homeostasis in different organs including the cardiovascular system. In patients with diabetes, endothelial dysfunction appears to be a consistent finding and was shown to be responsible for cardiovascular diseases including atherosclerosis, diabetic cardiomyopathy, diabetic retinopathy and diabetic kidney disease. Hyperglycemia, a hallmark of diabetes causes induction of a plethora of pathological pathways leading to endothelial dysfunction. Several studies have reported endothelial senescence in patients with diabetes ^1,2^ and cultured cells that were exposed to high glucose ^3,4^. Indeed, such senescent phenotype is a hallmark of cardiovascular anomalies in diabetes setting both *in vitro* and *in vivo*.

Senescence particularly leads to a phenotype known as senescence associated secretory phenotype. Two types of senescence can be accounted based on the environment and subsequent activity related with cell injury and division of the cell ^5^. The first is replicative senescence wherein a cell after dividing for many cycles can no longer propagate majorly because of excessive DNA damage leading to suppression of cell cycle mediators responsible for checkpoints at G1/S phase. The second is damage induced senescence in which stress due to toxic mediators or injury based inflammation leads to curtailing of cell division. In this case, the cell experiences a total arrest but is metabolically active ^6^. In such cases, these stressed cells start secreting various chemokines like TNF-α, IL-6 which lure immune cells such as macrophages and neutrophils towards the site of assault. This occurs preferentially through high amount of ROS activity by the mitochondria. The NF-κB transcriptional pathway along with the cGAS-STING pathway is primarily involved as p65 acts as a mediator for many of the signaling pathways progressing towards senescence ^7^. Interestingly, p65 known to interact with TERF2IP, part of shelterin complex, in promoting endothelial senescence. According to recent report, TERF2IP was found to shuttle from the nucleus to the cytoplasm causing p65 phosphorylation enabling it to translocate to the nucleus where it acts as a transcriptional inducer for endothelial senescence.

Shelterin complex is important as it helps in protecting chromosomal ends from damage thereby regulating cellular senescence. The complex composed of six proteins, namely-TPP1, TERF2IP, TRF1, TRF2, POT1, and TIN2. TRF1 and TRF2 are responsible for recruiting TERF2IP to form a complex by forming a D-loop at the telomeric ends ^8^. Loss of these proteins seems to trigger senescence either by hampering cell division or through telomere shortening. In vascular inflammation, shelterin knockouts have been found to progress to senescence, whereas if these were induced for a gain in expression, it was found to reduce inflammation. However, in a clinical setting, the difference could only be seen in high grade inflammation, particularly in lymphocytes^9^. For senescence to proceed, it would involve expression of the cell cycle control proteins – CDK 40, p16, p21, p27, particularly the tumor suppressor gene p53. A normally dividing cell would stop proliferation under two conditions; the first being replicative senescence and the second induced due to stress or deterioration, in this case inflammation ^10^. Although endothelial senescence was described in hyperglycemia setting, any role of shelterin protein in such phenotype have never been reported. We previously reported a H3K27me3 dependent repression of *KLF2* and *KLF4* genes resulting in endothelial inflammation ^11^. We also described hyperglycemia-dependent induction of partial mesenchyme-like phenotype of EC by elevating H3K4me3 level via MLL2-WDR82 protein of SET1/COMPASS complex which supported Jagged1 and Jagged2 expression and Notch activation ^12^. In this study, we report an increase in the expression of TERF2IP and TRF2 in EC upon exposure to intermittent high glucose. We also show that these proteins regulate the expression of classic cellular senescence associated proteins, p53 and p21, thereby causing senescence in EC upon hyperglycemia challenge. Interestingly, the expression of TERF2IP and TRF2 is also regulated by H3K4me3. This is a significant finding as no other study has reported the epigenetic regulation of TERF2IP and TRF2.

## Material and Methods

### Cell culture

Human umbilical vein endothelial cells (HUVEC) at passage 2 were purchased from Hi Media (CL002, Mumbai, India) and cultured using HiEndoXLTM Endothelial Cell Expansion medium (#AL517, Hi Media) supplemented with 2% endothelial growth supplement, 5% fetal bovine serum (FBS), and 1% pen/strep. HUVEC were grown in HiEndo XL cell medium with a basal glucose level of 5.5 mM. EA.hy926, an immortalized human umbilical vein cell, were purchased from ATCC (#CRL-2922, ATCC) and used as human EC in this study. The cells were cultured and passaged every 2 or 3 days in Dulbecco’s modified Eagle’s medium (DMEM) (Hi Media Laboratories) supplemented with 10% FBS (Hi Media Laboratories) and 1% penicillin/streptomycin (Sigma-Aldrich, St. Louis, MO, USA). Both cell types were cultured at 37°C and in a humidified incubator with a 5% CO2 atmosphere. After achieving a 70% confluency, HUVEC were then subjected to intermittent time kinetics of hyperglycemia, alternating between normal glucose level (5.5 mM) and high glucose level (25 mMan referred to as intermittent hyperglycemia and alternating between 12 h of high glucose (25 mM) and 12 h of normal glucose (5.5 mM) cycle for 3 cycles, totaling 72 h of treatment time^11^. EA.hy 926 cells were made tolerant to 15mM glucose through multiple passages by lowering down glucose concentration from normal growth condition of 25mM. The resulting cell line was then exposed to 30mM of glucose to achieve equivalent intermittent high glucose treatment condition as HUVEC while 15mM served as basal glycemic condition.

### Animal Dissection and Treatment Conditions

All experimental procedures for the rat aorta studies were approved by the Institutional Animal Ethics Committee of BITS Pilani, Pilani Campus, under the approval number IAEC/RES/27/12/Rev-1/13/17. Male Wistar rats aged 10 to 14 weeks were selected for the *ex vivo* experiment. The animals were provided with a normal chow diet. The animals were then sacrificed and in incision was made from the ventral end to excise the heart. Phosphate buffered saline (PBS) was used to perfuse the heart and the aorta to remove any blood cells ^13^. The primary aortas were then collected and the fatty tissue layers were removed, followed by cutting into cylindrical pieces measuring 2 mm in length to obtain aortic rings. These rings were washed with PBS and further cultured in HiEndo endothelial cell growth medium in 24-well plates for the experiment. The aortic rings were subjected to an intermittent high glucose (25mM) treatment, as explained earlier ^11^, along with treatment of OICR 9429 (#SML1209; Sigma Aldrich; 10μM)^13^. On completion of treatment, the tissue fractions were homogenized, suspended in RIPA lysis buffer (#9806, Cell Signaling Technology), and sonicated. Protein estimation was done by Bradford assay, followed by SDS-PAGE and immunoblot. Antibodies used for probing are summarized in the table (Table 1).

### Inhibitor Treatment Condition and RNA Silencing in Cultured Endothelial Cells

Prevalidated human-specific MLL2 small-interfering RNA (Silencer® MLL2 siRNA #AM16708) and control siRNA (SignalSilence® Control siRNA#6568) were purchased from Cell Signaling Technology. TERF2IP siRNA and TRF2 siRNA were designed and ordered from GeneCust (Boynes France). MLL2 siRNA was ordered from Thermo Fisher. The concentration of siRNA used was 40 nM. OptiMEM (#31985070, Thermo Fisher Scientific,Waltham, MA, USA) and lipofectamine 2000 (#11668030, Thermo Fisher Scientific,Waltham, MA, USA) was used to transfect the HUVEC cells with the specific siRNA. The cells were initially kept in the transfection medium for 4 hours after which the medium was replaced with HiEndo endothelial culture medium, which was later subjected to intermittent hyperglycemia treatment ^11^.

For EA.hy926, DMEM (#AL294A, HiMedia, Mumbai, India) culture medium was for culturing. EA.hy926 cells that normally grows at 25 mM glucose containing DMEM were initially acclimatized to grow in DMEM conating 15mM glucose. These 15mM adapted cells were then exposed to 30mM glucose containing DMEM to achieve intermittent high glucose treatment condition. Upon treatment, cells were finally harvested upon completion of the intermittent high glucose treatment condition, as specified earlier. For comparative analysis, all non siRNA - transfected cells were transfected with scrambled siRNA. For all MLL2 siRNA inhibition studies, an MLL-WDR5 inhibitor OICR9429 was used at a final concentration of 10μM. DMSO was used as vehicle control.

### Subcellular Fractionation

HUVEC exposed to intermittent hyperglycemia were harvested and washed with PBS. Upon centrifugation, the cell pellet was re-suspended in ice cold PBS containing 0.1% Nonidet P-40 (NP-40) and 1% protease inhibitors (#P8340, Sigma-Aldrich, MI, USA). The samples were then centrifuged for 10 min at 10,000 rpm and the nuclear fraction was obtained as pellet, whereas the supernatant was collected as the cytoplasmic fraction. Furthermore, the nuclear lysate was obtained by resuspending the nuclear pellet in PBS containing 0.1% Nonidet P-40 (NP-40) and 1% protease inhibitor (#P8340, Sigma). Both the nuclear and cellular fractions were sonicated for 10 seconds of two cycles each with a holding period of 5seconds to cool down and then separated by SDS-PAGE and immunoblotted for TRF2 and TERF2IP. GAPDH and histone H3 were used as loading controls for cytoplasmic and nuclear localization respectively. All details about the antibodies are provided in Table 1.

### Co-Immunoprecipitation

Proteins were extracted from HUVEC using RIPA lysis buffer (#9806, Cell Signaling Technology) containing protease inhibitor (#P8340, Sigma-Aldrich). 1000ug of total protein was used for pulldown with antibody for TERF2IP and p65. Antibodies specific to MLL2, p65, and TERF2IP from CST were procured from CST, and a dilution of 1:1000 was used for the study. Resulting bands were visualized by Chemi-Doc using the Clarity™ (#1705061, Bio-Rad Laboratories) or Clarity™ Max (#1705062, Bio-Rad Laboratories) Western Blotting ECL Substrates and analyzed by expression levels relative to 5% input obtained using ImageJ software. All details about the antibodies are provided in Table 1.

### Immunofluorescence Imaging and Analysis

HUVEC were grown on gelatin (#TC041, HiMedia) coated coverslips up to 70% confluency. After completion of treatment, they were washed with PBS and fixed with 4% ice-cold paraformaldehyde. Cells were incubated with 0.1% Triton X for permeabilization^12^. Cells were then incubated with BSA (1%) for 1 h, followed by overnight incubation with TERF2IP antibody or TRF2 antibody (1:1000). Cells were then washed with PBS to remove unbound primary antibody, and were subsequently incubated with Alexa fluor 555 secondary antibody (1:4000; # A32732, Thermo Fisher Scientific) for 2 h. (1:5000; # R415, Thermo Fisher Scientific) for 30 min, followed by incubation with DAPI (# D9542, Sigma-Aldrich) for nuclear staining. Fluorescence images were captured using a Zeiss ApoTome.2 microscope (Carl Zeiss, Jena, Germany), and intensities were measured using ImageJ software. For Co-immunofluorescence, a similar procedure was performed to stain the EA. hy926 cells as described earlier. In these experiments, cells were incubated with MLL2 antibody (1:1000) overnight. Next day, cells were washed with PBS to remove unbound primary antibody and incubated with anti-rabbit F(ab’)2 secondary antibody tagged with Alexa Fluor 488 for 2 h (1:2000, #4412S, CST). Cells were then again incubated with p65 primary antibody (1:1000) overnight and successively incubated with secondary antibody tagged with Alexa Fluor IgG 555 (#A32732, Thermo Fisher Scientific) as per abovementioned protocol. After staining with DAPI (#D9542; Sigma-Aldrich) and mounting with 70% glycerol, they were imaged to find co-localization through LSM 880 Zeiss confocal microscopy. Analysis was performed using Image J software^11^.

### RNA Isolation

RNA isolation was performed following the manufacturer’s protocol (#15596026, TRIzol™ Reagent, Life Technologies, Thermo Fisher Scientific). HUVEC were grown in six-well plates up to 70% confluency and subjected to the intermittent hyperglycemia treatment regime. After 72 h, treated cells were incubated in Trizol reagent. Upon collecting the cells in Trizol reagent, organic layer separation was carried out using chloroform, and further RNA in the aqueous layer was precipitated using isopropanol. Next, precipitated RNA was washed with 75% ethanol, and RNA pellets were air dried. Finally, RNA pellets were dissolved in sterile nuclease-free water, and quantity and quality were analyzed through NanoDrop measurement^11^.

### cDNA Synthesis and Quantitative Analysis through Reverse Transcriptase-Quantitative Polymerase Chain Reaction

rReverse transcriptase quantitative polymerase chain reaction (RT-qPCR) was performed to quantify the different genes at the transcript level. In brief, total RNA (1 µg) was taken from the cDNA preparation using iScript™ cDNA Synthesis Kit (#1708891, Bio-Rad Laboratories, Hercules, CA, USA). Prior to cDNA synthesis, isolated RNAs were pre-incubated with DNAse to remove any DNA contamination. Real-time PCR was then performed using iTaq™ Universal SYBR® Green Supermix (#1725124, Bio-Rad Laboratories), with a total master mix volume of 10 µL. Analysis was carried out by calculating delta-delta Ct. GAPDH was used as the housekeeping gene. All the sequence of the primers for TERF2IP, TRF2, TRF1, TPP1 and GAPDH are provided in Table 2.

### Chromatin Immunoprecipitation (ChIP) and Subsequent Quantitative PCR

A ChIP assay was performed using an Imprint® Chromatin Immunoprecipitation Kit (#86652 CUT & RUN assay kit, Cell signaling Technology). In brief, exponentially growing HUVEC (70% cell density) were subjected to intermittent high glucose treatment. Treated HUVEC were harvested (1*10^6^ cells), washed, and cross-linked with 1% formaldehyde in HiEndoXL Endothelial Cell Expansion medium (10 min at room temperature). After washing in PBS, the cell pellet was re-suspended and tagged with Concanavalin A Beads. Pulldown was done by H3K4me3 antibody at a dilution of 1mg/ml according to manufacturer’s protocol. Only input samples was sheared by sonication for 30 s pulses. The samples were then washed, reverse cross-linked, and treated with proteinase K to obtain purified DNA fragments through silica column. qPCR was performed using primers targeted to amplify regions of human gene promoters (Table 3).

### Cell cycle analysis using propidium iodide staining and flow cytometry

EA.hy926 cells seeded at 60% density were subjected to a five day treatment regime of intermittent high glucose and a combination of intermittent high glucose with TERF2IP siRNA or TRF2 siRNA. All were used at a concentration of 40 nM. OptiMEM containing lipofectamine 2000 (#11668030, Thermo Fisher Scientific,Waltham, MA, USA) and siRNA were subjected to HUVEC for 4 h. OptiMEM was later replaced with DMEM culture medium and subjected to the intermittent high glucose (30 mM) treatment condition. Cells were finally harvested upon completion of the intermittent high glucose treatment condition, as specified earlier^11^. For comparative analysis, all non siRNA-transfected cells were transfected with scrambled siRNA. Cells were harvested by trypsinization and washed with PBS followed by 5000 rpm for 10 min at 4°C, the pellet was re-suspended in a fixative (100 μL of PBS and 900 μL of ice cold 70% ethanol) and incubated at room temperature for 45 minutes. Cells were centrifuged at 5000rpm and the pellet was re-suspended in 400 μL PBS with 50 μL of propidium iodide (#P4170, Sigma-Aldrich) staining solution (PI; 2 mg/ml). The samples were then incubated in dark for 30 min followed by acquisition using flow cytometer (Cytoflex, Beckmann Coulter) and analysis using CytExpert software.

### Immunoblotting

HUVEC were grown up to 70% confluency for high glucose treatment conditions. Medium was removed and cells were briefly washed with sterile 1X phosphate-buffered saline (PBS). RIPA buffer containing protease inhibitor was then used for protein extraction. Cells were incubated in RIPA buffer for 1 h, followed by sonication. Cell lysates were centrifuged (10,000 rpm for 8 min), followed by collecting the supernatant and measuring protein concentration using Bradford reagent. Equal amounts of protein for different treatment groups were then applied in SDS-PAGE, followed by transferring the proteins onto a nitrocellulose membrane. Membranes were then blocked in 5% nonfat milk or 5% bovine serum albumin for 1 h. Membranes were then probed overnight at 4°C with different primary antibodies. The blots were then incubated with a peroxidase-conjugated anti-rabbit or anti-mouse IgG antibody (1:2000) (# 7074 or #7076, respectively; Cell Signaling Technology). The antibody–antigen reactions were detected using the Clarity™ or Clarity™ MaxWestern Blotting ECL Substrates (Bio-Rad). Densitometry analysis was performed using ImageJ software^11–13^. All details of the antibodies used for the study are detailed in Table 1.

### Flow cytometric analysis of apoptotic cells using Annexin V-propidium iodide (PI) staining

Dead Cell Apoptosis Kit with Annexin V FITC and PI for flow cytometry (#V13242, Thermo Fisher Scientific) was used to study the effects of intermittent high glucose exposure on endothelial apoptosis. To perform this, cells were seeded in 24 well dishes at a density of 1×10^5^ cells/well. The cells were challenged with intermittent high glucose. Next, treated cells were harvested and washed twice in cold PBS. Following centrifugation, the cell pellet was resuspended in 1X annexin-binding buffer (approx. 1×10^6^ cell/ml), and 5 µl of FITC annexin V (component A) and 1 µl of PI (100 µg/ml) were added to each 100 µl of cell suspension. After 15 min incubation at room temperature, the cell suspension was further diluted with 400 µl PBS and the stained cells were analyzed using the CytoFLEX flow cytometer (Beckman Coulter). Apoptosis, both early and late, was determined from the respective Annexin V and Annexin V-PI positive quadrants, after subtracting the autofluorescence. Acquired data were analyzed using CytExpert software ^12,13^.

### Statistical analysis

All the values are expressed as the mean ± SD. All analysis data in bar graphs are presented as relative to control treatment condition. Statistical significance was determined by one-way ANOVA followed by Tukey’s Post Hoc test for multiple groups comparison or by two-tailed Student’s *t* test for comparison between two groups, unless otherwise stated. Statistical analyses were performed using GraphPad Prism software. A p value of less than 0.05 was considered statistically significant.

## Results

### Intermittent high glucose selectively elevated the level of p53 and p21 to cause endothelial senescence without causing apoptosis

Induction of ROS upon hyperglycemia exposure was described to be responsible for cellular senescence ^14^. eNOS-mediated nitric oxide production caused inhibition of endothelial senescence upon hyperglycemia challenge ^15^. We previously reported robust reduction in eNOS transcript and protein levels thereby prompting us to study the effect of intermittent hyperglycemia on senescence ^11^. Furthermore, we also reported no alteration in two proliferation markers Cyclin D3 and c-Myc in EC exposed to hyperglycemia ^12^. Exit from cell cycle and pause in cell proliferation is generally the end point of cellular senescence. Therefore, we evaluated the changes in cyclin inhibitors like p53, p21, p16 and damage induced senescence marker lamin B1. Protein expression of both p53 (Figure 1A,G) and p21 (Figure 1B,F) increased significantly upon intermittent hyperglycemia challenge indicating hyperglycemia did lead to changes in cell proliferation. No robust change was observed in expression of other senescence markers such as p16 (Figure 1C), TNF-α (Figure 1D), lamin B1 and MMP3 (Figure 1E). We also confirmed the increase in p53 level in intermittent hyperglycemia-exposed rat aorta (Figure 1H). As p53 and p21 were well established to halt cell cycle at G1/S checkpoint and our previous findings revealed no alteration in level of Cyclin D3 and cMyc ^12^, we wondered that intermittent hyperglycemia is leading to cell cycle arrest at G0/G1 phase. To test this, we performed cell cycle analysis using PI staining followed by flow cytometric analysis to determine the total DNA content of each cell. In doing so, we detected significant number of intermittent hyperglycemia exposed EC arrested at G0/G1 phase than normal glucose treated group (Figure 1I). Moreover, five days of intermittent hyperglycemia exposure caused a significant increase in cells having G0/G1 arrest (Figure 1I). Because cellular senescence is responsible for cell apoptosis, we therefore detected the relative fraction of apoptotic cells after intermittent hyperglycemia exposure. No changes in percentage of apoptotic cell (Figure 1J) was detected when EC were challenged with intermittent hyperglycemia, thus disclosing absence of senescence-dependent endothelial apoptosis phenotype in the present experimental settings. To exclude the possibility that the intermittent hyperglycemia effect was due to osmolality changes, we simultaneously analyzed these senescence markers in HUVEC exposed to intermittent high mannitol, which generates a comparable osmolality that is similar to intermittent hyperglycemia. In so doing, we detected no changes in p53 and p21 level in EC exposed to IHM (Figure S1A).

**Figure 1.**
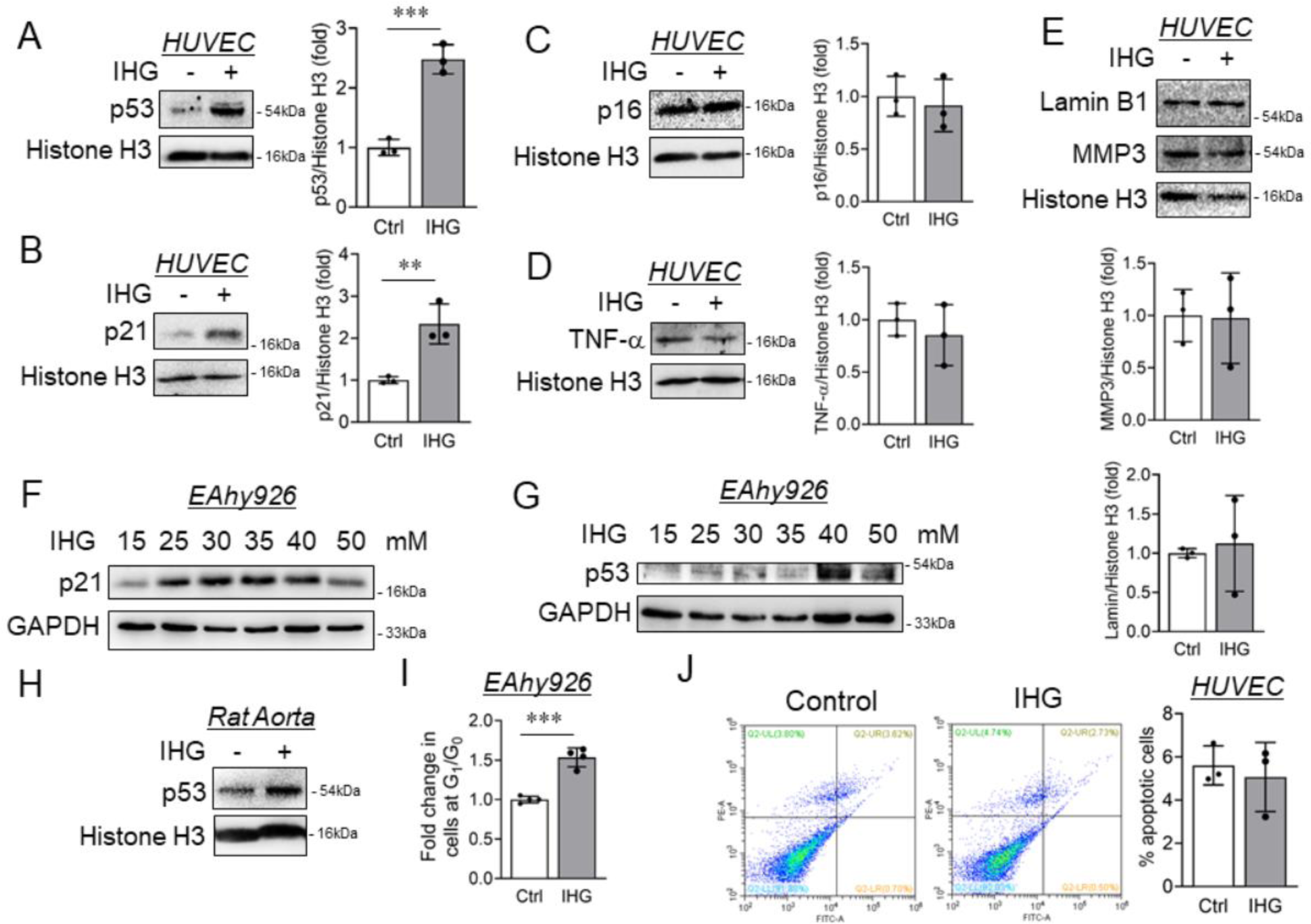
Intermittent high glucose exposure enhanced endothelial senescence without altering apoptosis by inducing selective expression of p53 and p21. (A-E) Immunoblot analysis of HUVEC exposed to intermittent high glucose to detect the level of p53 (A), p21 (B), p16 (C), TNF-α (D), Lamin B1 and MMP3 (E) (n=3). (F-G) Level of p21 (F) and p53 (G) were detected in EA.hy926 challenged with intermittent high glucose (n=3). Densitometric quantification of the blots were represented. (H) Immunoblot analysis of p53 in tissue lysate collected from rat aorta exposed to intermittent high glucose *ex vivo* (n=3). (I) Measuring the level of apoptosis in HUVEC treated with intermittent high glucose using Annexin V-PI staining followed by flow cytometry analysis; distribution plots of early and late apoptotic cells (E) and quantified data that include both early and late apoptotic cell population. (n=3). (J) Cell cycle analysis of intermittent high glucose challenged EA.hy926 cells using PI staining followed by flow cytometry analysis to quantify cells at G_1_/G_0_, S-phase and G_2_/M phase (n=4). Data represented as fold change relative to control (normal glucose cells) for cells at G_1_/G_0_ phase. Values represent the mean ± SD. **P < 0.01, and ***P < 0.001, by unpaired t test.

### Intermittent high glucose elevated the level of p65 and Shelterin complex proteins TERF2IP and TRF2

Oxidative stress and inflammation disrupt shelterin in vascular cells causing telomere dysfunction and cellular senescence ^9^. Moreover, deletion of shelterin complex protein like TERF2IP inhibited d-flow-induced EC senescence, apoptosis, and activation ^16^. Hence, we evaluated the expression level of proteins related to shelterin complex in EC challenged with intermittent hyperglycemia condition. Transcript and protein level expression analysis revealed significant increase in TERF2IP (Figure 2A,E,M) and TRF2 (Figure 2B,F,L) in EC exposed to intermittent hyperglycemia. In contrast, the level of other two components of the shelterin complex; TRF1 (Figure 2C,G) and TPP1 (Figure 2D,H) remain to be unaltered upon intermittent hyperglycemia challenge. Analysis of TERF2IP (Figure S1B) and TRF2 (Figure S1A) in intermittent high mannitol challenged EC exhibited no alteration in these proteins, thus excluding any contribution of osmolality in intermittent hyperglycemia dependent increase in TERF2IP and TRF2 levels.

**Figure 2.**
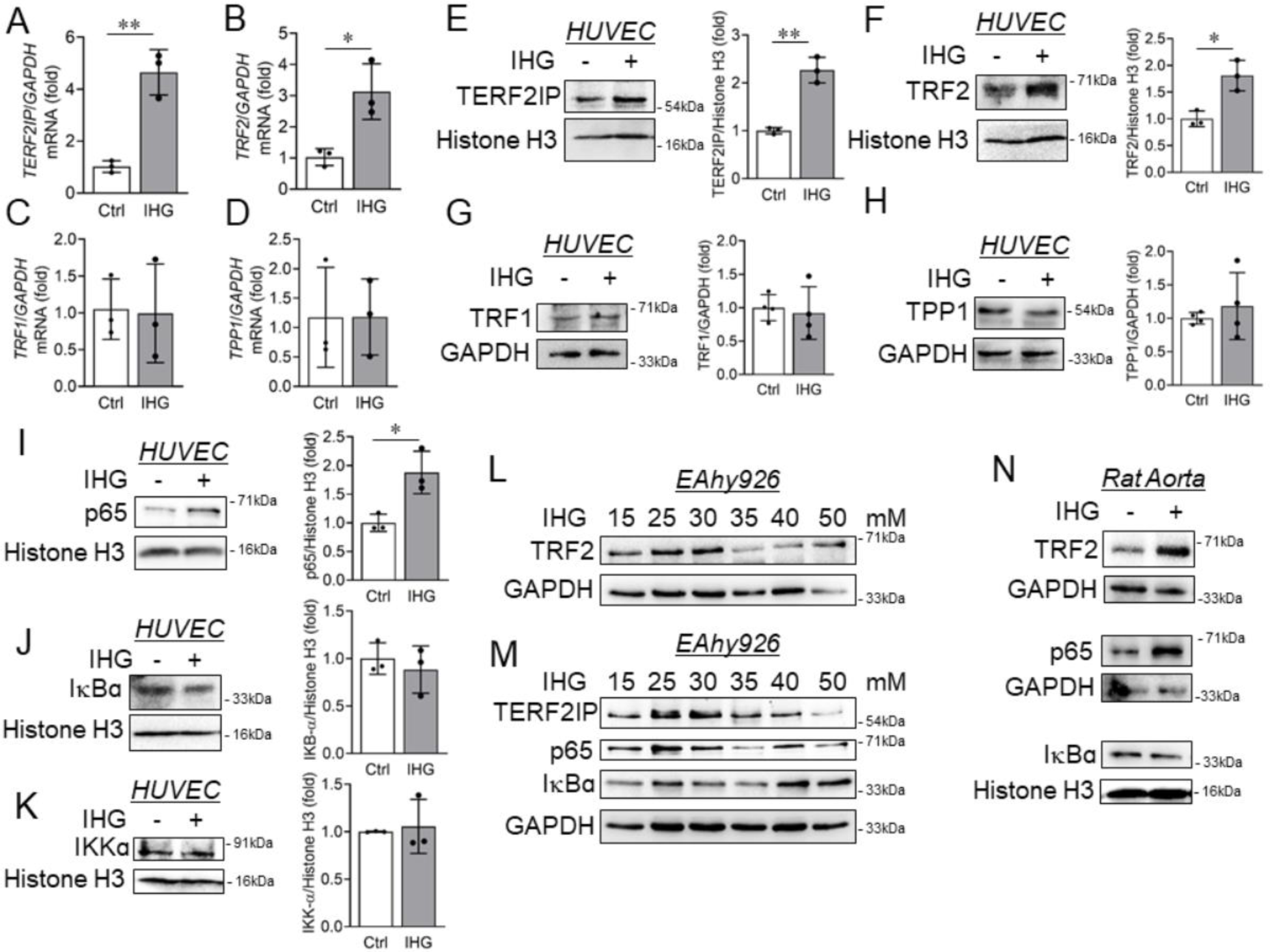
Shelterin proteins TERF2IP-TRF2 and p65 are elevated in EC exposed to intermittent high glucose. (A-D) Transcript-level expression of *TERF2IP* (A), *TRF2* (B), *TRF1* (C) and *TPP1* (D) measured through RT-qPCR technique in HUVEC challenged with an intermittent high-glucose condition (n=3). (E-H) Immunoblot analysis of HUVEC exposed to intermittent high glucose to detect the level of TERF2IP (E), TRF2 (F), TRF1 (G) and TPP1 (H) (n=3). Densitometric quantification of the blots were represented. (I-K) The level of p65 (I), IκBα (J) and IKKα (K) were quantified in HUVEC challenged with intermittent high glucose (n=3). Densitometric quantification of the blots were represented. (L-M) Level of TRF2 (L), TERF2IP (M), p65 (M), and IκBα (M) (G) were detected in EA.hy926 challenged with intermittent high glucose (n=3). (N) Immunoblot analysis of TRF2, p65 and IκBα in tissue lysate collected from rat aorta exposed to intermittent high glucose *ex vivo* (n=3). Values represent the mean ± SD. *P < 0.05, and **P < 0.01, by unpaired t test.

The canonical NF-κB pathway has been shown to occur in many chronic diseases including obesity, inflammation, cardiovascular damage and type 2 diabetes. In addition, NF-κB signaling plays a pivotal role in endothelial senescence ^17^. p65 is an important transcription factor as it activates the downstream genes responsible for cell cycle arrest. Hyperglycemia and supported senescence mechanisms having p65 involvement as a mediator are widely known. We therefore analyzed the expression level of proteins involved in NF-κB analysis. We found a profound increase in expression of p65 in intermittent hyperglycemia as opposed to untreated samples (Figure 2I,M). Expression of IκB-α (Figure 2J,M) and IKK-α (Figure 2K) remain unchanged in EC treated with intermittent hyperglycemia condition. Therefore, induction of p65 expression upon intermittent hyperglycemia exposure in EC was likely to be independent of IκB-α and IKK-α action on p65 activity. Intermittent high mannitol exposure to EC did not alter the level of p65 (Figure S1B) suggesting osmolality changes was not responsible for increase in p65 level upon intermittent hyperglycemia exposure.

To validate these findings in more relevant *in vivo* model, we detected the changes in these genes in isolated rat aortas which are exposed to intermittent high glucose treatment condition *ex vivo*. We detected a significant increase in level of TRF2 and p65 (Figure 2N) in tissue lysates of aortas that undergone intermittent hyperglycemia treatment condition while no alteration in IκB-α (Figure 2N) was detected in these tissue.

### Increased level of H3K4me3 upon intermittent high glucose epigenetically regulated the expression of TERF2IP, TRF2 and p65

Because we observed an increased level of TERF2IP, TRF2 and p65, we therefore decided to decipher the underlying mechanism responsible for such selective increase in gene expression. We previously showed that intermittent high glucose led to increase in H3K4 trimethylation in EC through preferential increase in MLL2 and WDR82 level ^12^. With this in mind, we decided to understand the role of H3K4me3 in intermittent hyperglycemia dependent increase in TERF2IP, TRF2 and p65 level. ChIP-qPCR analysis revealed heightened enrichment of H3K4me3 in the promoter region of *p65* (Figure 3A), *TERF2IP* (Figure 3B) and *TRF2* (Figure 3C) genes.

**Figure 3.**
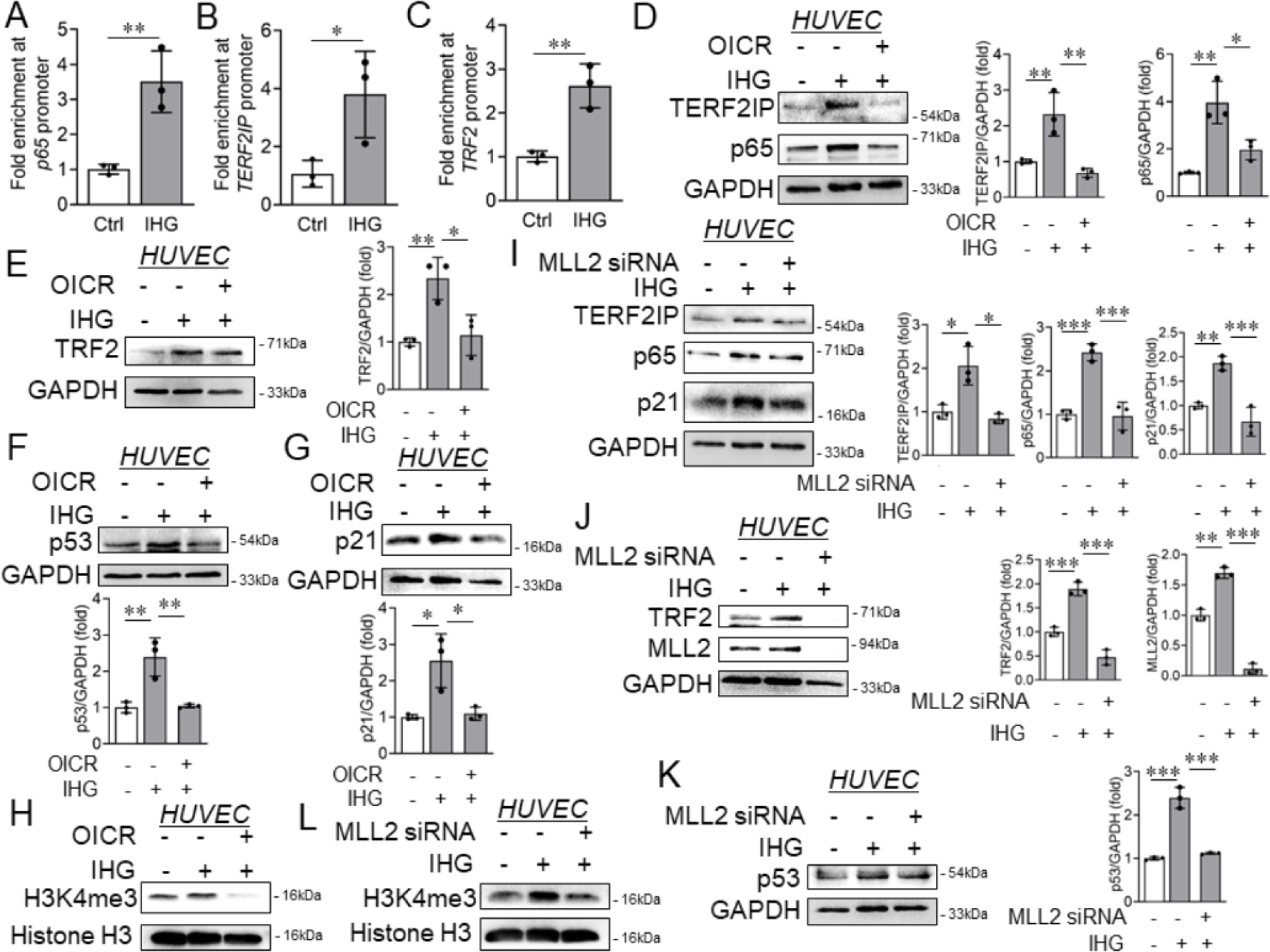
Heightened H3K4me3 was responsible for intermittent high glucose dependent increase in p65, TERF2IP and TRF2 levels. (A-C) ChIP-qPCR for *p65* (A), *TERF2IP* (B) and *TRF2* (C) gene promoters following enrichment with H3K4me3 antibody in HUVEC challenged with intermittent high glucose treatment condition (n = 3). (D-H) Immunoblot analysis of HUVEC lysate collected from cells pre-treated with OICR-9429 (10 μM) followed by exposing to intermittent high glucose. Immunoblots were probed with antibodies against TERF2IP and p65 (D), TRF2 (E), p53 (F), p21 (G) and H3K4me3 (H) proteins/modifications along with densitometry quantification of the blots (n=3). (I-L) Immunoblot analysis of HUVEC lysates collected from cells transfected with MLL2 specific siRNA followed by challenging with intermittent high glucose and probed for TERF2IP, p65, and p21 (I), TRF2 and MLL2 (J), p53 (K), and H3K4me3 (L) proteins/modifications along with densitometry quantification of the blots (n=3). Values represent the mean ± SD. *P < 0.05, **P < 0.01, and ***P < 0.001, by Tukey’s multiple comparisons test of one-way ANOVA.

To further ascertain that elevated promoter level enrichment of H3K4me3 was the reason of enhanced expression of TERF2IP, TRF2 and p65, we further analyzed the expression of these genes in EC incubated with selective pharmacological inhibitors of SET1/COMPASS methylation complex or specific siRNA to MLL2. Selective inhibition of the activity of SET1/COMPASS complex neutralized the intermittent hyperglycemia-driven enhanced expression of TERF2IP (Figure 3D), TRF2 (Figure 3E) and p65 (Figure 3D). In parallel, we detected that selective knockdown of MLL2 reversed intermittent hyperglycemia dependent increase in expression of TERF2IP (Figure 3I), TRF2 (Figure 3J) and p65 (Figure 3I). Because we suspected that elevated level of p65, TERF2IP and TRF2 was responsible for enhanced level of senescence associated biochemical changes, we therefore detected the level of p53 and p21 in EC treated with selective inhibitors of SET1/COMPASS complex activity or MLL2 specific siRNA and exposed to intermittent hyperglycemia. In so doing, we reported attenuation of intermittent hyperglycemia induced increase in the expression of p21 (Figure 3G,I) and p53 (Figure 3F,K) in EC. As we reported previously ^12^, we confirm the reduction in the level of H3K4me3 in EC that were exposed to intermittent hyperglycemia in combination with selective inhibitors of SET1/COMPASS complex activity or MLL2 specific siRNA (Figure 3H,L). In addition, we also showed no changes in the level of H3K4me3 in EC treated with intermittent high mannitol condition thereby eliminating the role of osmolality in the catalysis of H3K4me3 (Figure S1C).

### p65 in conjunction with MLL2 was responsible for intermittent high glucose-dependent increase in TERF2IP and TRF2

A relatively recent study indicated TERF2IP is able to translocate from nucleus to cytoplasm during endothelial inflammation and associate with p65 to cause activation of NF-κB signaling in EC ^18^. Therefore, we studied the effect of intermittent high glucose exposure on the subcellular localization of TERF2IP and TRF2. In so doing, we found both TERF2IP (Figure S2A) and TRF2 (Figure S2B) were primarily localized in the nucleus while no changes in the subcellular localization of these proteins were detected between control and intermittent hyperglycemia-treated cells. To further validate, we performed a cell fractionation assay followed by immunoblot analysis. Although a significant increase in both nuclear and cytosolic localization in TERF2IP was detected due to an elevation in total TERF2IP level, relative distribution remained constant (Figure S2C). In contrast, TRF2 did not show any changes in sub-cellular localization (Figure S2D). The association of TERF2IP with p65 is well documented ^16^, we thus performed a co-immunoprecipitation study to evaluate TERF2IP association with p65 upon intermittent hyperglycemia challenge. Interestingly, as reported earlier in EC, we detected an association of p65 with TERF2IP, however, a modest reduction in the association of TERF2IP and p65 was detected upon intermittent high glucose exposure (Figure S2E).

We next investigated the role of p65 in the regulation of intermittent hyperglycemia-dependent induction of TERF2IP and TRF2. We next inhibited p65 using the pharmacological inhibitor, JSH-23 and evaluated its effect on intermittent high glucose-induced TERF2IP and TRF2 expression. Inhibition of NF-κB activation by JSH-23 completely neutralized intermittent hyperglycemia-dependent elevation in TERF2IP (Figure 4A) and TRF2 level (Figure 4B). Moreover, inhibition of p65 also reversed intermittent hyperglycemia-supported increase in p21 (Figure 4C) and p53 (Figure 4D) levels.

**Figure 4.**
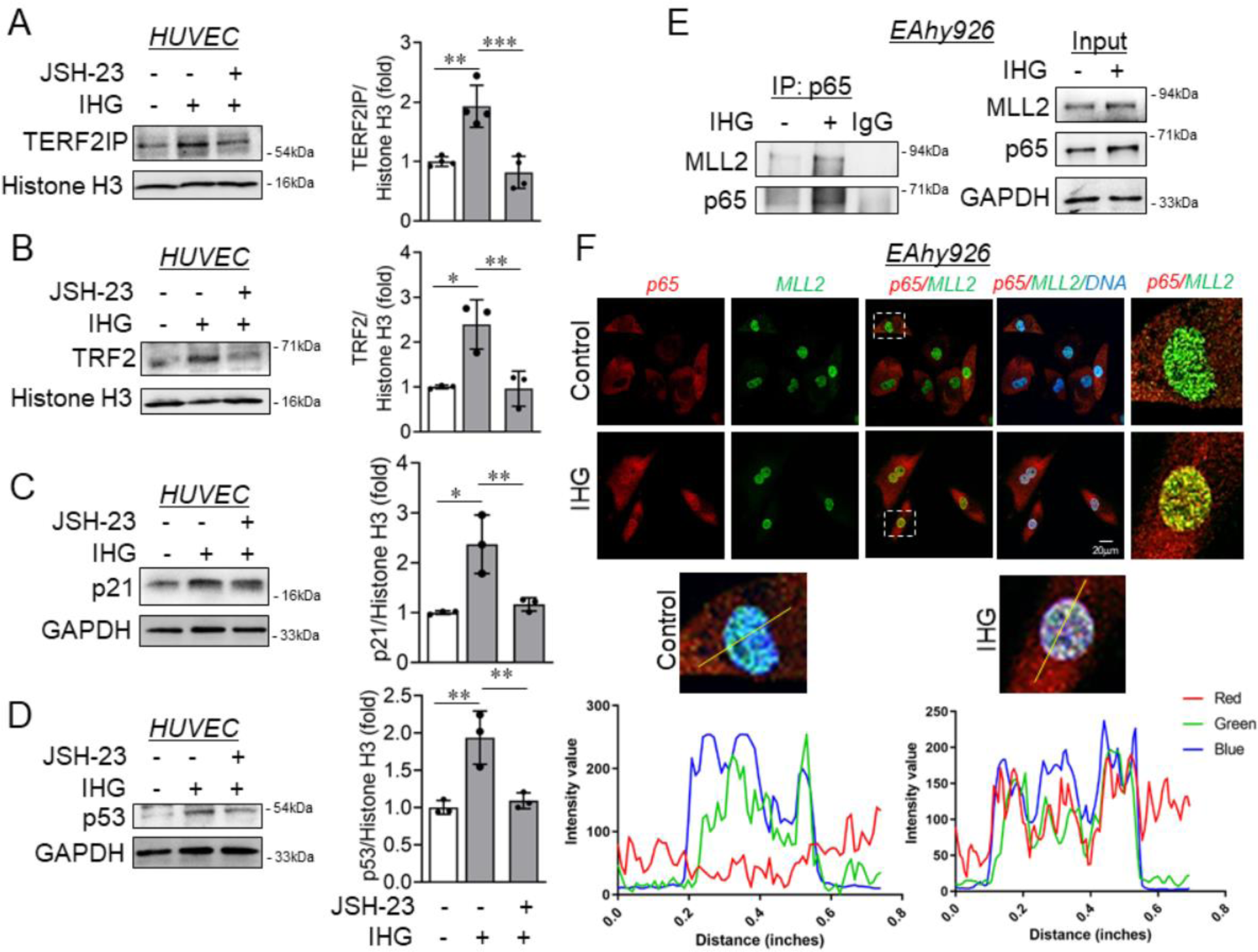
p65 in conjunction with MLL2 was responsible for intermittent high glucose dependent increase in shelterin proteins TERF2IP-TRF2 level and cellular senescence markers p21-p53. (A-D) Immunoblot analysis of HUVEC lysate collected from cells pre-treated with JSH-23 (10 μM) followed by exposing to intermittent high glucose. Immunoblots were probed with antibodies against TERF2IP (A), TRF2 (B), p21 (C) and p21 (D) proteins along with densitometry quantification of the blots (n=3). (E) Co-immunoprecipitation with p65 antibody, followed by immunoblotting for MLL2, p65, and GAPDH in either immuno-precipitated or total cell lysate (Input) sample (n=3). (F) Immunofluorescence analysis and co-staining of EA.hy926 exposed to intermittent high glucose for p65 and MLL2. DAPI staining to visualize the nucleus is shown in blue (n = 3). Line graph indicating pixel-wise intensities of green (MLL2) and red (p65) fluorescence in the yellow line highlighted areas within the image. Values represent the mean ± SD. *P < 0.05, **P < 0.01, and ***P < 0.001, by Tukey’s multiple comparisons test of one-way ANOVA.

As per our findings, both inhibitions of MLL2 or p65 caused a reversal of intermittent hyperglycemia-induced TERF2IP and TRF2 expression, we next wondered whether MLL2 and p65 work in tandem with each other to regulate the expression of TERF2IP and TRF2. As MLL2 and p65 were never reported to associate with each other in any cell type, we questioned whether intermittent hyperglycemia can confer such an association between p65 and MLL2. Co-immunoprecipitation analysis confirmed the association of MLL2 and p65 in EC, moreover, such association is enhanced upon the intermittent high glucose challenge (Figure 4E). We confirmed such association of MLL2 and p65 in the nucleus of EC using immunofluorescence analysis (Figure 4F).

### Pharmacological inhibition of MLL2 reversed the level of TRF2, p65 and senescence marker p53 in intermittent hyperglycemia exposed rat aortic ring model ex vivo

Upon confirming MLL2 and p65 regulation of shelterin protein TEFR2-IP and TRF2 and further intermittent hyperglycemia effect on cellular senescence marker p53, we next assessed the effect of such inhibition in ex vivo model. Blocking SET/COMPASS complex activity using pharmacological inhibitor OICR-9429 in *ex vivo* model of rat aortic rings caused a reversal of intermittent hyperglycemia-induced expression of TRF2 (Figure 5A). Furthermore, such inhibition of H3K4me3 catalysis also neutralized intermittent high glucose-dependent increase in p65 level and cellular senescence marker p53 (Figure 5B).

**Figure 5.**
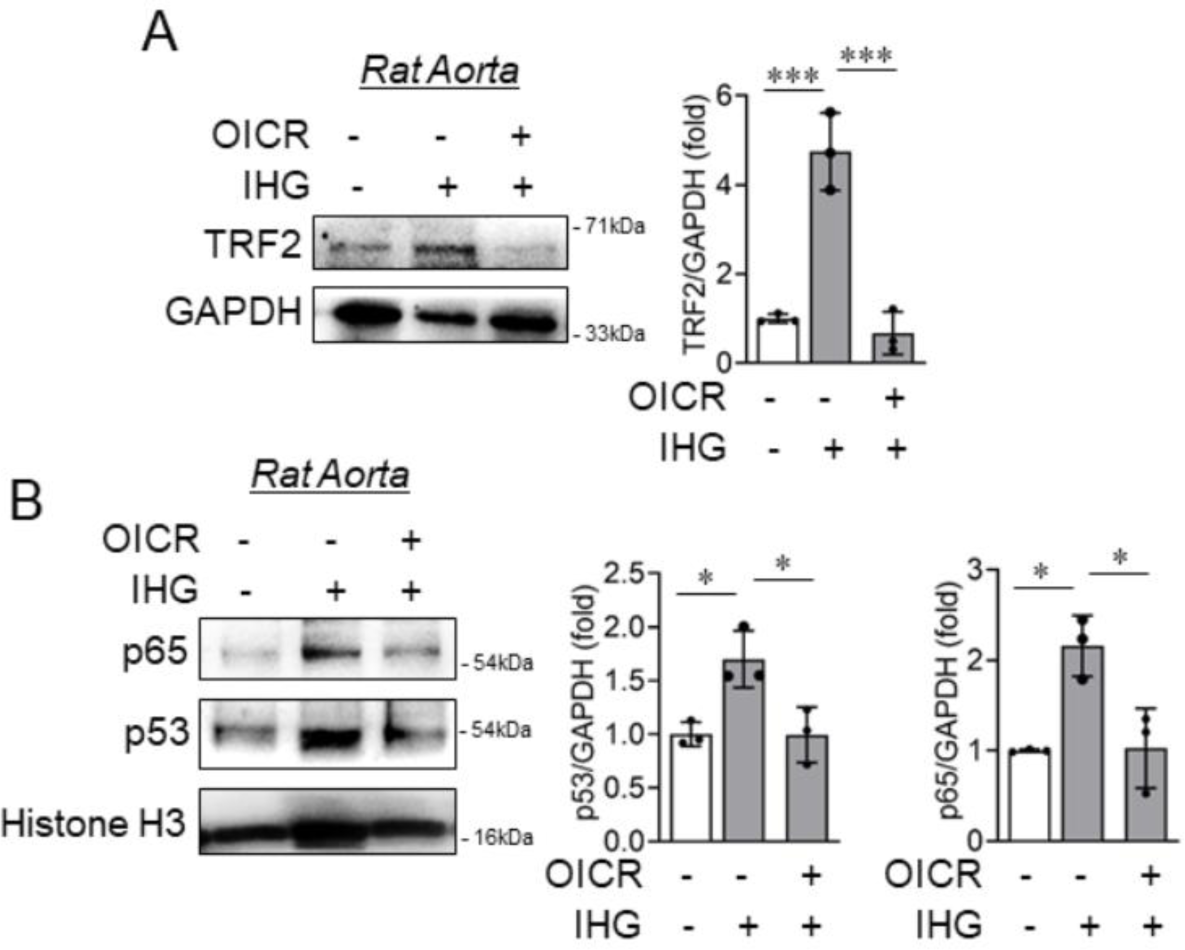
Catalytic inhibition of MLL blocked intermittent high-glucose-dependent increase in TRF2, p65 and p53 in rat aortic rings *ex vivo*. (A-B) Immunoblot analysis of tissue lysates collected from rat aorta treated with intermittent high glucose in combination with OICR-9429 (10μM) and probed for TRF2 (A), p65 (B), and p53 (B) along with densitometry quantification (n=3). Values represent the mean ± SD. *P < 0.05, and ***P < 0.001, by Tukey’s multiple comparisons test of one-way ANOVA.

### Knockdown of TERF2IP and TRF2 reversed p53 and p21 level and associated endothelial senescence imparted by intermittent high glucose

Induction of senescence state in cells is dependent on the relative expression of TERF2IP ^19^ and TRF2 ^20^. Through a recent study, aberrant expression of TERF2IP has been known to cause aging-related phenotype ^21^. Therefore, we wondered whether upregulation in the level of shelterin protein TERF2IP and TRF2 upon intermittent high glucose exposure caused changes in senescence-associated biochemical markers, p21 and p53. To this end, we transfected cells with TERF2IP siRNA or TRF2 siRNA and exposed them to intermittent hyperglycemia treatment condition. TERF2IP knockdown as confirmed through immunoblot (Figure 6A) completely reversed intermittent high glucose-induced increase in the expression of p21 (Figure 6A) and p53 (Figure 6B). In parallel, TRF2 knockdown (Figure 6C) in EC overrode the changes observed in p21 (Figure 6C) and p53 (Figure 6D) level upon intermittent high glucose challenge. Knockdown of TERF2IP or TRF2 also completely abrogated intermittent high glucose induced cell cycle arrest of EC at G1/G0 phase (Figure 6E).

**Figure 6.**
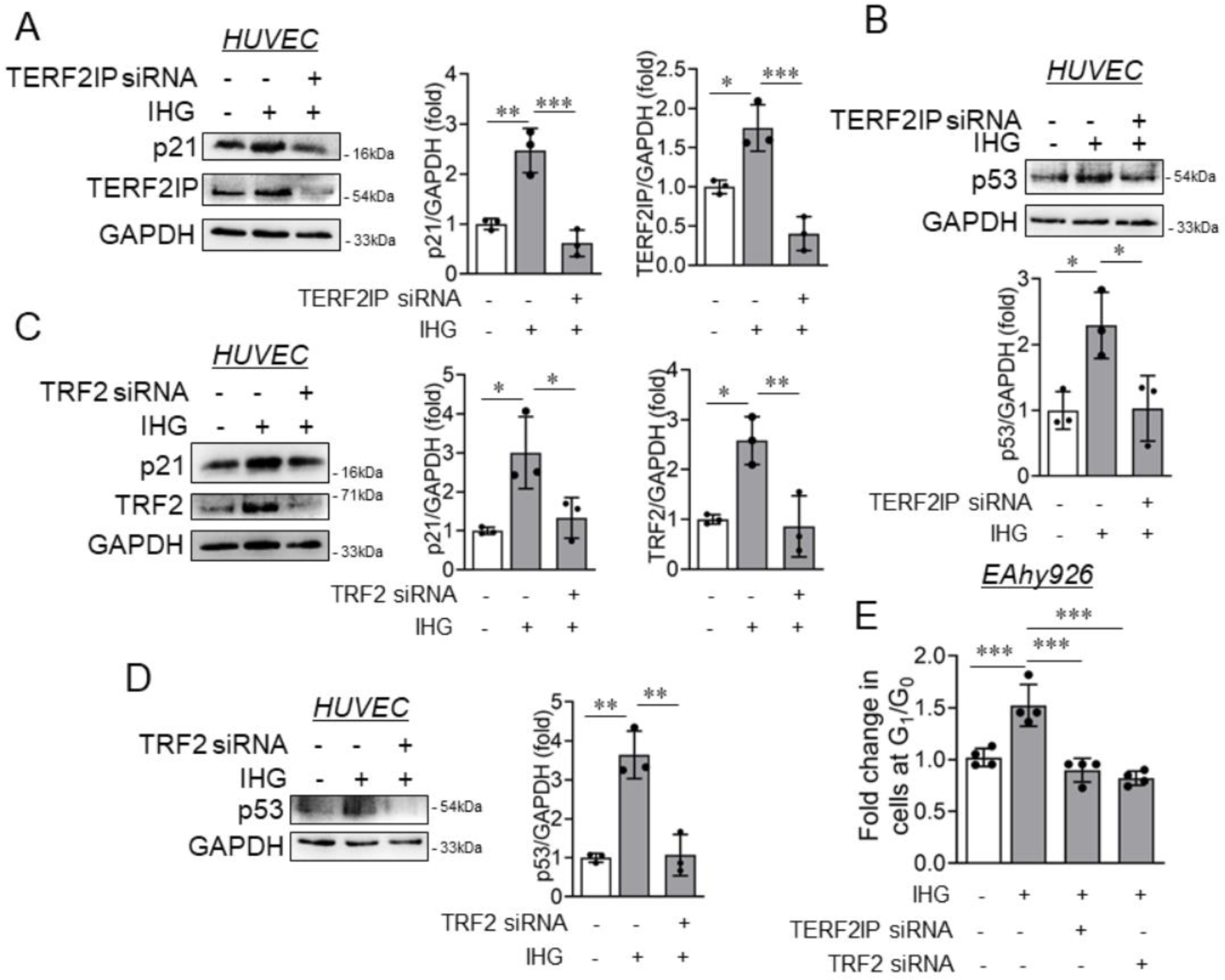
Knockdown of either TERF2IP or TRF2 reversed p53 and p21 level including endothelial senescence imparted by intermittent high glucose exposure. (A-B) Immunoblot analysis of HUVEC lysates collected from cells transfected with MLL2 specific siRNA followed by challenging with intermittent high glucose and probed for p21 (A), TERF2IP (A) and p53 (B). Densitometry quantification of the blots were plotted as graph (n=3). (C-D) HUVEC transfected with TRF2 specific siRNA were challenged with intermitted high glucose treatment condition followed immunoblot analysis with the cell lysate for p21 (C), TRF2 (C), and p53 (D) along with densitometry quantification of the blots (n=3). (E) Cell cycle analysis of TERF2IP or TRF2 knockdown and intermittent high glucose challenged EA.hy926 cells using PI staining followed by flow cytometry analysis to quantify cells at G_1_/G_0_ (n=4). Data represented as fold change relative to control (normal glucose incubated cells) for cells at G_1_/G_0_ phase. Values represent the mean ± SD. *P < 0.05, **P < 0.01, and ***P < 0.001, by Tukey’s multiple comparisons test of one-way ANOVA.

## Discussion

Senescence can occur as a result of DNA damage, cytokine stress or replicative dysfunction. Moreover, senescence has been known to cause inflammation through induction of ROS and increase in NADPH oxidases ^22,23^. Such phenomenon is dependent on replication of progeny through induction of cell cycle or be a pseudo enhancement damage induced senescence. Transcriptional regulation through histone modifications like H3K9me3, H3K4me3 or H3K27me3 at specific foci can also act to promote senescence in cells. However, many such mechanisms of regulation are studied based on progression of senescence towards inflammation. Furthermore, this is not always the case as senescence activated inflammatory cells is a longterm response and because of chronic manifestation of such phenotype, these cells could no longer be senescent in nature. Cells existing in senescence or growth arrest are more detrimental as they can no longer maintain a constant metabolic state. Through the present study we describe the effect of intermittent high glucose on induction of a senescent response in endothelial cells through transcriptomic regulation of p53 and p21 by two shelterin proteins TERF2IP and TRF2. We report that intermittent high glucose causes MLL2-H3K4me3-mediated transcriptional regulation of p65 which in turn binds and recruits MLL2 to cause enhanced transcriptional activity of TERF2IP and TRF2 genes. Targeting either NF-κB or MLL2 reversed high glucose induced expression of TERF2IP and TRF2, further reverting the senescence-associated changes in biochemical markers such p53 and p21. Moreover, knockdown of either TERF2IP or TRF2 abrogated hyperglycemia-dependent increase in p53 and p21 levels including reversal of G1/G0 cell cycle arrest of the EC (Figure 7).

**Figure 7.**
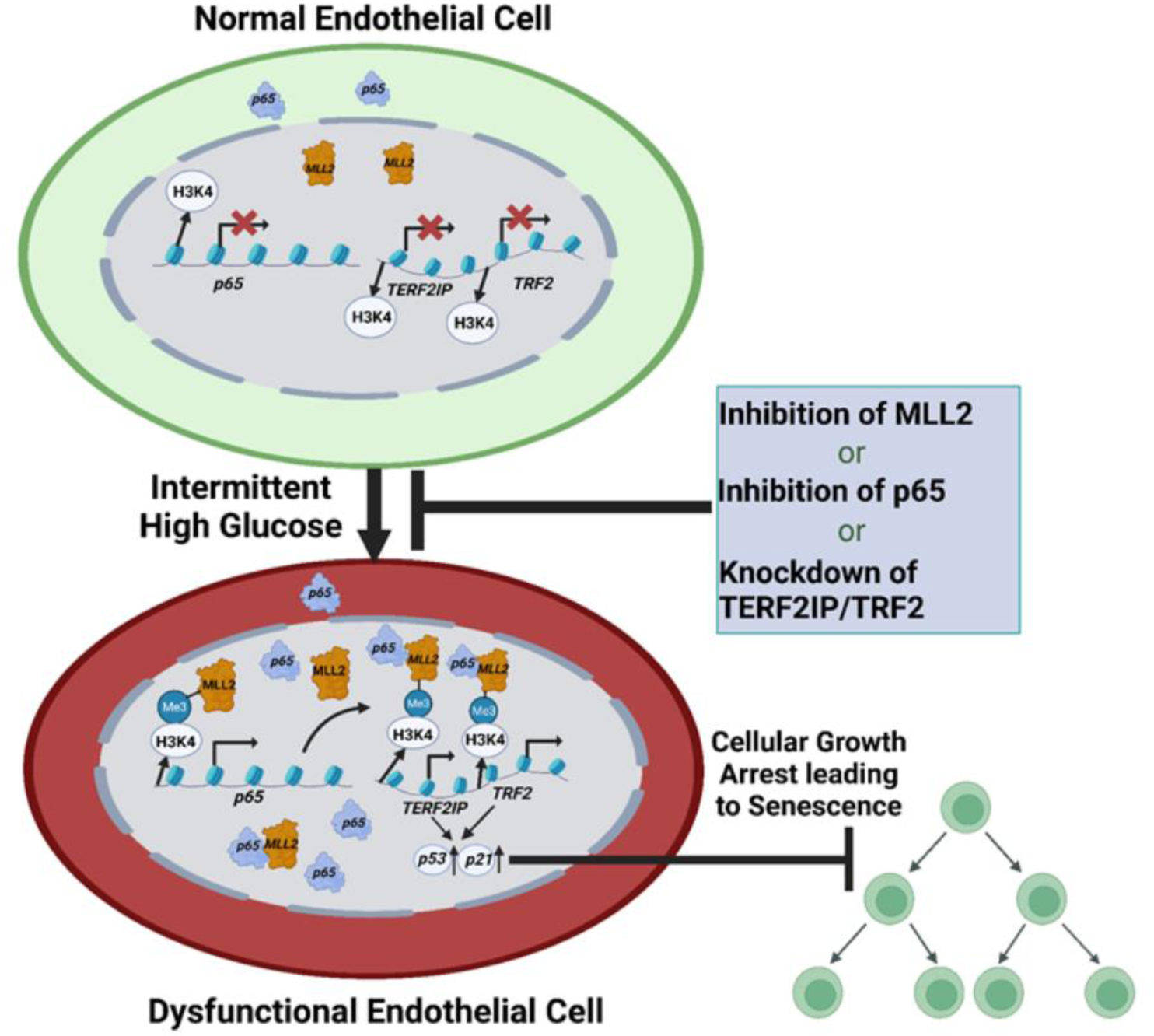
Schematic illustration of the intermittent high-glucose-driven induction of TERF2IP and TRF2 through H3K4me3-p65 pathway to cause endothelial senescence. In normal glucose condition, limited level of p65 and MLL2 exist within endothelial cells, thereby supporting minimum expression of shelterin associated genes TERF2IP and TRF2. In such condition, a basal level of cellular senescence associated genes p21 and p53 are expressed, thus maintaining normal endothelial turn-over. Upon intermittent high-glucose challenge, endothelial cells exhibit a heightened level of H3K4me3 by MLL2 induction supporting transcriptional derepression of p65 gene and increasing its relative abundance. Further, MLL2 in conjunction with p65 caused H3K4me3 dependent elevation in TERF2IP and TRF2 level which supported the expression of cellular senescence associated genes p53 and p21 leading to endothelial senescence upon hyperglycemia shock. In contrast targeting either MLL2 or p65 or TERF2IP or TRF2 either by pharmacological inhibitor or siRNA reversed intermittent high glucose dependent increase in p21 and p53 level and cellular senescence. The schematic diagram was prepared using BioRender (Agreement number: EX25CEZQD).

Shelterin proteins are responsible for preventing telomeric damage leading to telomeric shortening, further causing associated phenotype ^24^. However, in many reports, it has been observed that cells experience senescence even though they have long telomeres, and differing levels of TERF2IP and TRF2 proteins ^18,25,26^. Indeed, TERF2IP and TRF2 were shown to have non telomeric activity and can respond to oxidative stress in many cells like fibroblasts, mesenchymal cells, smooth muscle cells, skin cells^18,27,28^. A telomeric independent pathway has been shown in which both TERF2IP and TRF2 can act as transcription factors in angiogenesis, maintaining pluripotency in stem cells ^29,30^. Both can also act independently as opposed to telomeric capping in which TRF2 recruits TERF2IP and binds to telomeric ends in a D loop. TERF2IP knockout mice were shown to be incapable of atheroma progression while ApoE knockout mice when introduced with TRF2 were shown to have a greater amount of senescence in liver ^31^. However, in humans, senescence is inconsistent and is different in various organs as they age. Both mice and rats have longer telomeres than human, yet they are more likely to manifest senescence phenotype due to fast developmental maturity decline ^32^. In such cases, senescence also depends on the stage at which the cell is in and its current environment. Therefore, the presence of shelterin complex and associated proteins to stabilize the telomere could not be the sole determinant for senescence or growth arrest phenotype of different cells. In addition, these studies clearly indicated shelterin independent functions of TERF2IP/TRF2 including working as a transcriptional regulator or causing induction of cellular senescence when overexpressed. In parallel, we observed that induction of TERF2IP and/or TRF2 in a hyperglycemia settings caused induction of cellular senescence/growth arrest supported by enhanced level of TERF2IP and/or TRF2. Furthermore, gene specific knockdown of TERF2IP or TRF2 reversed high glucose-dependent endothelial senescence/growth arrest indicating heightened expression of these genes are likely responsible for the manifestation of growth arrest phenotype.

p65 subunit of the NF-κB pathway is a transcriptional modulator and has been shown to regulate both p53 and p21 expression which are key part of cellular senescence pathways ^33^. Published work by Wang et al 2009, embryonic fibroblasts isolated from p65 knockout mouse showed to escape senescence at a faster rate than embryonic fibroblasts collected from p65 overexpressing mice ^34^. Such observation indicated a key role of p65 in imparting cellular senescence response. In an independent study forced expression of c-Rel, a member of the NF-κB transcription factor family was sufficient to induce senescence in fibroblasts by an oxidative-stress-related mechanism ^34^. Furthermore, p65 has also been shown to activate cell cycle genes responsible for G1-phase entry during cell division ^35^. We observed that intermittent hyperglycemia led to an increase in the p65 subunit with no difference in levels of IKK-α. Because we observed heightened level of shelterin proteins, TERF2IP and TRF2, we were curious to understand the crosstalk of TERF2IP/TRF2 and p65 pathways. TERF2IP in humans is responsible for inhibiting phosphorylation of the p65 subunit of NF-κB, through IKK-α making p65 competent for transcriptional activation ^18^. Such activation of p65 by TERF2IP was dependent on cytosolic translocation of TERF2IP to associate with p65 for further transactivation of the NF-κB signaling. However, our data did not indicate changes in the relative abundance of TERF2IP in cytosolic fraction of EC upon hyperglycemia exposure. Although, we detected association of TERF2IP with p65, such association remained unaltered upon hyperglycemia challenge. In contrast, nuclear masking of p65 by JSH-23 led to a repression in protein levels of both TERF2IP and TRF2 in endothelial cells in a hyperglycemia setting.

Many global histone modifications particularly H3K9me3 and H3K27me3 which lead to repression of Ras signaling, DNA damage response pathways, eventually lead to a stress induced premature senescence. The epigenetic regulator SET1/COMPASS particularly the MLL complex has been implicated in senescence. For instance, targeting MLL1 as a therapeutic strategy to deal with both oncogene induced senescence and senescence activated secretory phenotype as it activates the cell cycle genes namely p16 and p27. In another study, H3K4 methylation through SETD1A caused mitotic switch of cells and DNA-damage responses through regulation of different genes. Indeed, depletion of SETD1A caused chromosome misalignment and segregation defects including induction of senescent phenotype by transcriptional suppression of SKP2, which degrades p27 and p21 ^36^. Although SET/COMPASS proteins or H3K4me3 have been shown to repress the expression of senescence associated genes in cancer microenvironment, however, our study in hyperglycemia settings with EC strongly indicated MLL2 and its catalytic product H3K4me3 caused heightened expression of p65 and shelterin protein TERF2IP-TRF2 which in turn induced the expression of p53 and p21. Downregulation of the H3K4me3 mark via pharmacological inhibition or gene knockdown of MLL2 reversed the expression of p65, TERF2IP, TRF2, p53 and p21 including countermanding hyperglycemia-driven cellular senescence.

To summarize, our study revealed the role of TERF2IP and TRF2 in controlling the expression of cellular senescence associated genes p53 and p21 further causing endothelial senescence upon hyperglycemia challenge. This in overall been governed by the regulation of H3K4me3 and in downstream p65 when EC were exposed to high glucose conditions. Telomeric erosion and uncapping is a major cause for concern encompassing senescence, however in many studies, telomere length has been shown to be unchanged. Regardless, independent to its function in telomere stability, shelterin proteins have also been observed to act as transcription factors and regulators of protein activity such as p65 activation, thus ruling out a probability of telomere shortening as the only driver of shelterin regulation dependent cellular senescence response in hyperglycemia settings. Also, TERF2IP and TRF2 have been known to regulate cell cycle proteins during genotoxic stress. Concomitantly, p53 and 21 were both found to be downregulated on knockdown of TERF2IP and TRF2 gene transcripts, suggesting that intermittent hyperglycemia promotes activation of TERF2IP and TRF2, thereby regulating levels of p21 and p53 and leading to senescence. Exploring the role of shelterin proteins in diabetes could allow greater understanding of their role in onset and progression of diabetes and associated complications including cardiorenal diseases.

## Acknowledgments

We gratefully acknowledge the technical assistance of Mr. Suman Kumar for using the Confocal facility of BITS Pilani, Pilani Campus.

## Sources of Funding

This work was supported by Competitive Research Grant from the Department of Biotechnology, Govt. of India (BT/PR33144/MED/30/2170/2019) to SM. This work was also partly supported by an Adhoc Research Project from Indian Council of Medical Research, Govt. of India (55/1/2019-BMS) to S.M. S.T. was supported by a graduate fellowship from BITS Pilani. Y.T.K. and N.P.T. are supported by a graduate fellowship from BITS Pilani. S.R. is supported by a graduate fellowship from Department of Science and Technology-Innovation in Science Pursuit for Inspired Research fellowship (DST/INSPIRE/03/2019/000582).

## Notes

### Competing Interest Statement

The authors have declared no competing interest.

### Summary of Updates

Added the supplemental data file.

