## Supplemental Data for "Regulation of shelterin proteins TERF2IP and TRF2 by H3K4me3-p65 axis drives hyperglycemia dependent endothelial senescence"

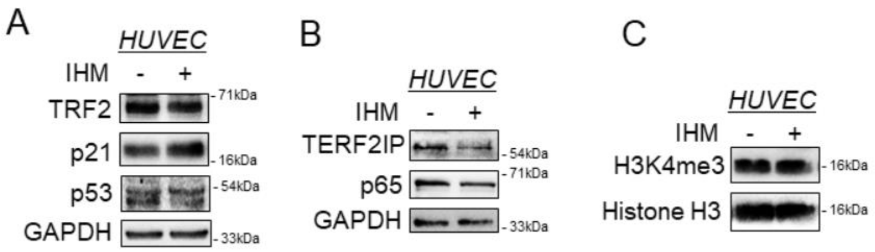


**Figure S1. Intermittent high mannitol exposure did not alter TERF2IP, TRF2, p65, p21, p53 and H3K4me3 level in HUVEC.** (A-C) Immunoblot analysis of HUVEC exposed to intermittent high mannitol (IHM) to detect the level of TRF2 (A), p21 (A), p53 (A), TERF2IP (B), p65(B), and H3K4me3 (C) (n=3).

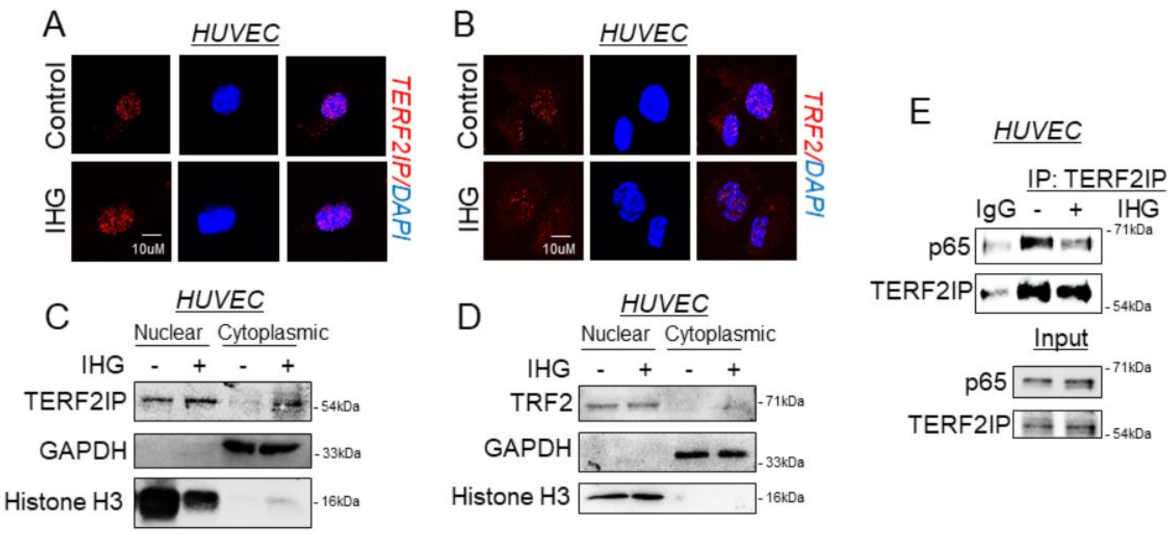


**Figure S2. Relative distribution of TERF2IP and TRF2 upon intermittent high glucose remain unaltered while p65 invariably associate with TERF2IP independent of intermittent high glucose challenge.** (A-B) Immunofluorescence analysis and co-staining of HUVEC exposed to intermittent high glucose for TERF2IP (A) and TRF2 (B). DAPI staining to visualize the nucleus is shown in blue (n=3). (C-D) Subcellular fractionation, immunoblotting, and quantitation of nuclear and cytosolic level of TERF2IP (C) and TRF2 (D) in intermittent high glucose-challenged HUVEC (n=3). (E) Co-immunoprecipitation with TERF2IP antibody, followed by immunoblotting for p65, and TERF2IP in either immuno-precipitated or total cell lysate (Input) sample (n=3).
